# PARP16 is a Druggable Regulator of Ribosome MARylation and Protein Homeostasis in Ovarian Cancer Cells

**DOI:** 10.64898/2026.04.01.715986

**Authors:** Sridevi Challa, Morgan Dasovich, Jonathan C. Abshier, Komal Pekhale, Lu Yang, Cristel V. Camacho, W. Lee Kraus

## Abstract

Cytosolic NAD□ synthesis supports ovarian cancer growth by enabling PARP16-dependent mono(ADP-ribosyl)ation (MARylation) of ribosomal proteins, thereby fine-tuning translation and maintaining protein homeostasis. While genetic depletion of PARP16 disrupts ribosome MARylation and impairs tumor cell growth, the therapeutic potential of pharmacologic PARP16 inhibition in this pathway remains unexplored. Here, we characterized the effects of DB008, a tool compound that functions as a selective inhibitor of PARP16, in ovarian cancer cells. Biochemical analyses demonstrated that PARP16 undergoes NAD□-dependent auto-MARylation and that NMNAT-2 supplies NAD□ to support this activity. DB008 potently inhibited PARP16 auto-MARylation in vitro. In ovarian cancer cells, DB008 engaged PARP16, reduced its MARylation, and decreased ribosome-associated MARylation. Consistent with PARP16 depletion, DB008 enhanced global protein synthesis, increased protein aggregation, and suppressed cell growth and anchorage-independent colony formation. CRISPR-mediated deletion of the *PARP16* gene in ovarian cancer cells abolished the effects of DB008 on translation, protein aggregation, and proliferation, demonstrating on-target activity. Moreover, cells expressing a PARP16 mutant resistant to DB008 were unaffected by inhibitor treatment, further confirming that the cellular effects of DB008 require on-target inhibition. Finally, DB008 significantly inhibited tumor growth in OVCAR3 xenografts, with on-target engagement of PARP16 in the xenograft tumors. Collectively, these findings establish PARP16 as a druggable regulator of ribosome MARylation and protein homeostasis in ovarian cancer and provide pharmacologic proof-of-concept that disrupting ribosomal MARylation impairs tumor growth.

## Introduction

Nicotinamide adenine dinucleotide (NAD□) is an essential metabolic cofactor that also serves as a substrate for ADP-ribosyltransferases, including members of the poly (ADP-ribose) polymerase (PARP) family (1–3). Because PARPs consume NAD□ during ADP-ribosylation reactions, their activity depends on NAD□ resynthesis through the salvage pathway, in which compartment-specific NMNAT enzymes generate NAD□ in distinct cellular locations (4, 5). The PARP family comprises 17 enzymes with diverse catalytic activities and subcellular localizations. While nuclear PARPs that catalyze poly(ADP-ribosyl)ation have been extensively studied, most PARP family members are cytosolic mono(ADP-ribosyl) transferases (MARTs) whose biological roles and therapeutic potential remain poorly defined (6–9). We recently demonstrated that cytosolic NAD□ synthesis supports ovarian cancer growth by enabling PARP16-dependent mono(ADP-ribosyl)ation (MARylation) of ribosomal proteins. PARP16-mediated ribosome MARylation fine-tunes translation by limiting polysome assembly, thereby maintaining protein homeostasis and preventing toxic protein aggregation (10). Genetic depletion of PARP16 disrupts ribosome MARylation, enhances translation, increases protein aggregation, and impairs ovarian cancer cell growth (10). These findings identify ribosomal MARylation as a previously unrecognized mechanism that couples NAD□ metabolism to translational control in cancer. However, whether this pathway is therapeutically targetable has remained unknown.

Rapidly proliferating cancer cells must maintain a precise balance between protein synthesis and protein quality control. Ribosomes serve as central hubs in this process, not only synthesizing proteins but also coordinating the recruitment of factors involved in folding and degradation (11–15). Post-translational modifications of ribosomal proteins—including phosphorylation, acetylation, ubiquitination, and other chemical modifications—have emerged as key regulators of translational output and ribosome specialization (16–18). Although ADP-ribosylation of translation factors is well characterized in the context of bacterial toxins that inactivate elongation factor 2 (E2F) and halt protein synthesis (19), endogenous ADP-ribosylation of ribosomal components as a regulatory mechanism in cancer has only recently begun to be appreciated.

In this study, we investigate the therapeutic potential of targeting PARP16-dependent ribosome MARylation in ovarian cancer. Recently, a tool compound that functions as a selective covalent inhibitor of PARP16, DB008, was described (20). DB008 was reported to have family-wide selectivity for PARP16 in vitro and covalently label PARP16 in cells, but whether DB008 mediates anti-cancer effects is unknown. Here, we characterize DB008 as a potent and selective inhibitor of PARP16 catalytic activity in ovarian cancer and demonstrate that pharmacologic inhibition phenocopies genetic loss of PARP16. DB008 suppresses ribosome MARylation, enhances protein synthesis, promotes protein aggregation, and impairs anchorage-independent growth of ovarian cancer cells in a PARP16-dependent manner. Importantly, DB008 inhibits tumor growth in vivo and engages PARP16 within xenograft tumors. Together, our findings provide pharmacologic proof-of-concept that disrupting cytosolic NAD□-dependent ribosome MARylation is a viable strategy to impair ovarian cancer growth.

## Results

### DB008 selectively and potently inhibits PARP16 activity

We recently demonstrated that PARP16-mediated MARylation of ribosomal proteins fine-tunes translation by limiting polysome assembly, thereby maintaining protein homeostasis and preventing toxic protein aggregation (10) (Fig. 1A). To explore whether this pathway is therapeutically targetable, we tested the recently developed selective PARP16 inhibitor compound DB008 (20). Using purified PARP16, we performed in vitro MARylation reactions and showed PARP16 autoMARylation in a NAD^+^-dependent manner (Fig. 1B). Based on the MARylation signals and NAD^+^ concentrations, we determined that the K_m_ of PARP16 for NAD^+^ was ∼200 µM (Fig. 1C), which is in agreement with previous literature (21, 22). One should note, however, that these measurements were made using a purified recombinant version of a PARP16 ex situ, removed from its native environment in the endoplasmic reticulum.

**Figure 1.**
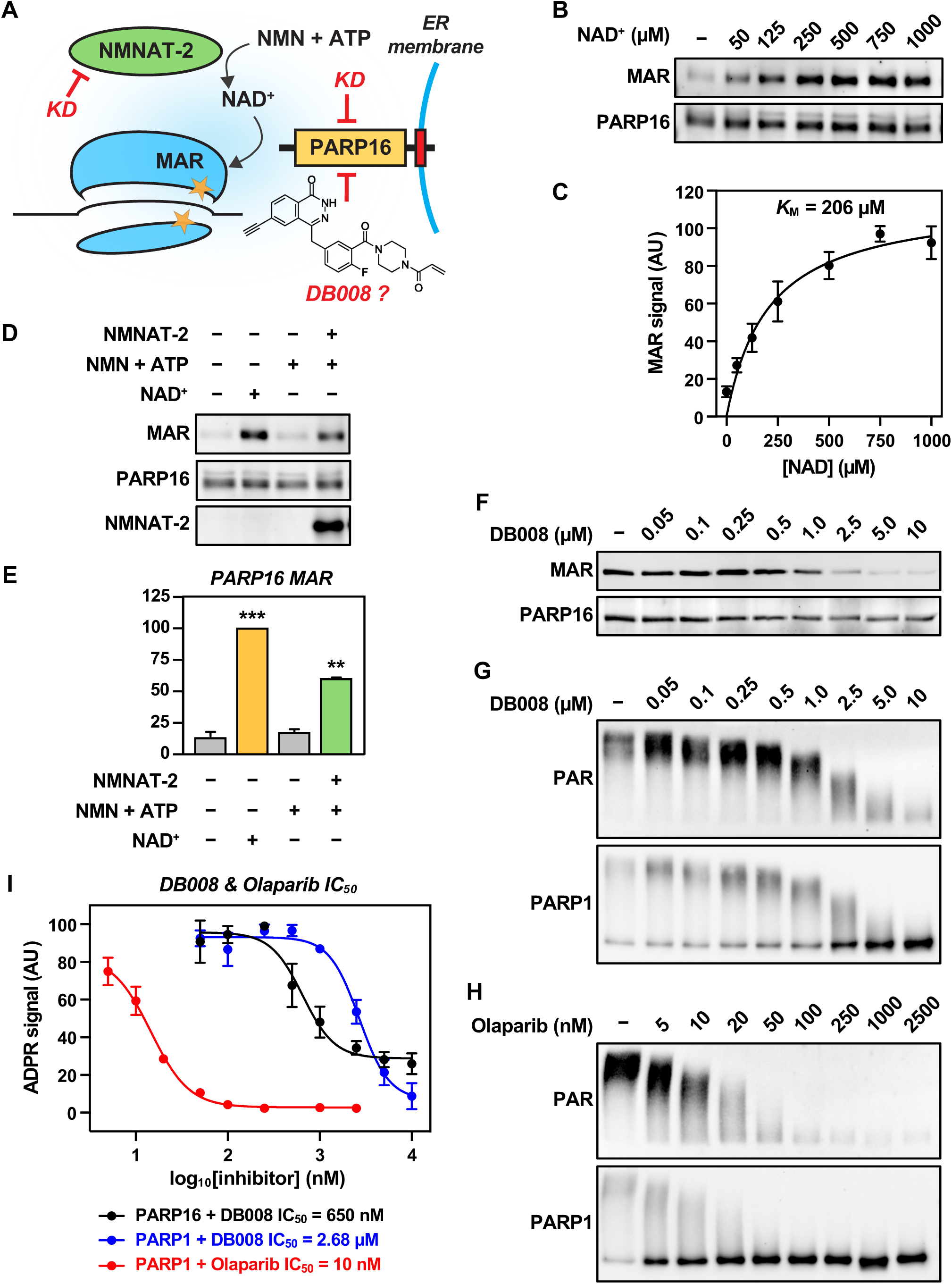
PARP16 biochemistry and its inhibition by DB008. **(A)** Schematic representation of NMNAT-2- and PARP16-dependent ribosome MARylation in ovarian cancer. **(B and C)** PARP16 MARylates itself in a NAD^+^-dependent manner. (B) Western blot of in vitro autoMARylation (MAR) reactions with PARP16 purified from *E. coli* in the presence of 0 – 1 mM NAD^+^. (C) MAR signals were plotted to determine the K_M_. Each point represents the mean ± SEM; n = 4. **(D and E)** NMNAT-2 can supply the NAD^+^ for PARP16 autoMARylation *in vitro*. (D) Western blot showing in vitro autoMARylation (MAR) reactions with PARP16 and NMNAT-2 purified from *E. coli* in the presence of 0.5 mM NMN/ATP or 0.5 mM NAD^+^. (E) Quantification of MAR signal from D. Each bar represents the mean + SEM; n = 2. Bars marked with asterisks are significantly different from the control; ANOVA; ** = p < 0.01, *** = p < 0.001. **(F)** DB008 inhibits PARP16. Western blot showing a DB008 dose-dependent reduction in PARP16 autoMARylation (MAR) *in vitro*. (**G and H**) Western blots showing PARP1 autoPARylation (PAR) in the presence of 0 – 10 µM DB008 (G) or 0 – 2.5 µM Olaparib (H). **(I)** DB008 is more potent against PARP16 than PARP1. Quantification of MAR and PAR signals from F–H. Each point represents the mean ± SEM; PARP16 & DB008, n = 4; PARP1 & DB008, n = 3; PARP1 & Olaparib, n = 2.

Next, we replaced the supply of NAD^+^ with purified NMNAT-2 and its substrates NMN and ATP. These in vitro reactions showed that NMNAT-2 can synthesize NAD^+^, which then serves as a substrate for PARP16, resulting in autoMARylation levels comparable to those observed with a direct supply of NAD^+^ (Figs. 1, D and E). Next, we tested the ability of DB008 to inhibit PARP16 activity in vitro and observed a dose-dependent reduction in PARP16 autoMARylation, with a significant reduction in signal when treated with 2.5 µM DB008 (Fig. 1F). To test selectivity, we repeated this in vitro assay using purified PARP1 with increasing doses of DB008 or olaparib as a control (Figs. 1, G and H). By measuring the levels and PARP16 autoMARylation and PARP1 autoPARylation, these results demonstrate that while DB008 has some inhibitory effect on PARP1 at higher doses, DB008 is more potent against PARP16 (IC_50_ 650 nM) compared to PARP1 (IC_50_ 2.7 µM) (Fig. 1I) Furthermore, DB008 is a covalent, irreversible inhibitor while inhibition of PARP1 is reversible. Overall, the results confirm that DB008 selectively and potently inhibits PARP16 activity.

### DB008 inhibits PARP16, decreases ribosome MARylation, and increases protein synthesis in ovarian cancer cells

Having verified the inhibitory effects of DB008 on PARP16 in vitro, we next tested the effect in cell-based assays. First, we examined the proteome-wide selectivity of DB008. OVCAR3 cells ectopically expressing FLAG-tagged PARP16 were treated with increasing doses of DB008. Taking advantage of the clickable alkyne handle, we incubated cell lysates with CuSO_4_ and TAMRA-azide and observed selective labeling of a band correlating in size with the molecular weight of FLAG-PARP16 (Fig. 2A), demonstrating proteome selectivity. We also immunoprecipitated FLAG-PARP16 from OVCAR3 lysates and showed autoMARylation activity of PARP16 in cells, which was potently inhibited by DB008 in a dose-dependent manner (Fig. 2B), consistent with in vitro assays. Using proximity ligation assay (PLA), we also demonstrated a significant PARP16-specifc loss of autoMARylation upon treatment with DB008 using a MAR-detection reagent and FLAG to detect ectopically expressed PARP16 (Fig. 2, C and E). Second, we evaluated the effect of DB008 on ribosome MARylation and protein translation. PLA verified RPS6-specific transMARylation, which was significantly reduced with DB008 treatment (Fig. 2, D and F). In line with this observation, the ribosome fraction of OVCAR3 cells had a significant decrease in MARylation signals with DB008 treatment compared to control (Fig. 2G). We also labeled the cells with puromycin after treatment with increasing doses of DB008. We observed a robust increase in puromycin incorporation (i.e., translation), along with increased cleaved caspase-3 signal (Figs. 2, H and I), corroborating our previous studies with genetic depletion of PARP16 (10). These findings demonstrate that DB008 effectively inhibits PARP16 activity in cells. Importantly, DB008 alters downstream biological outcomes, including reduced ribosome MARylation and increased protein synthesis—phenotypes we previously showed are detrimental to ovarian cancer cell survival.

**Figure 2.**
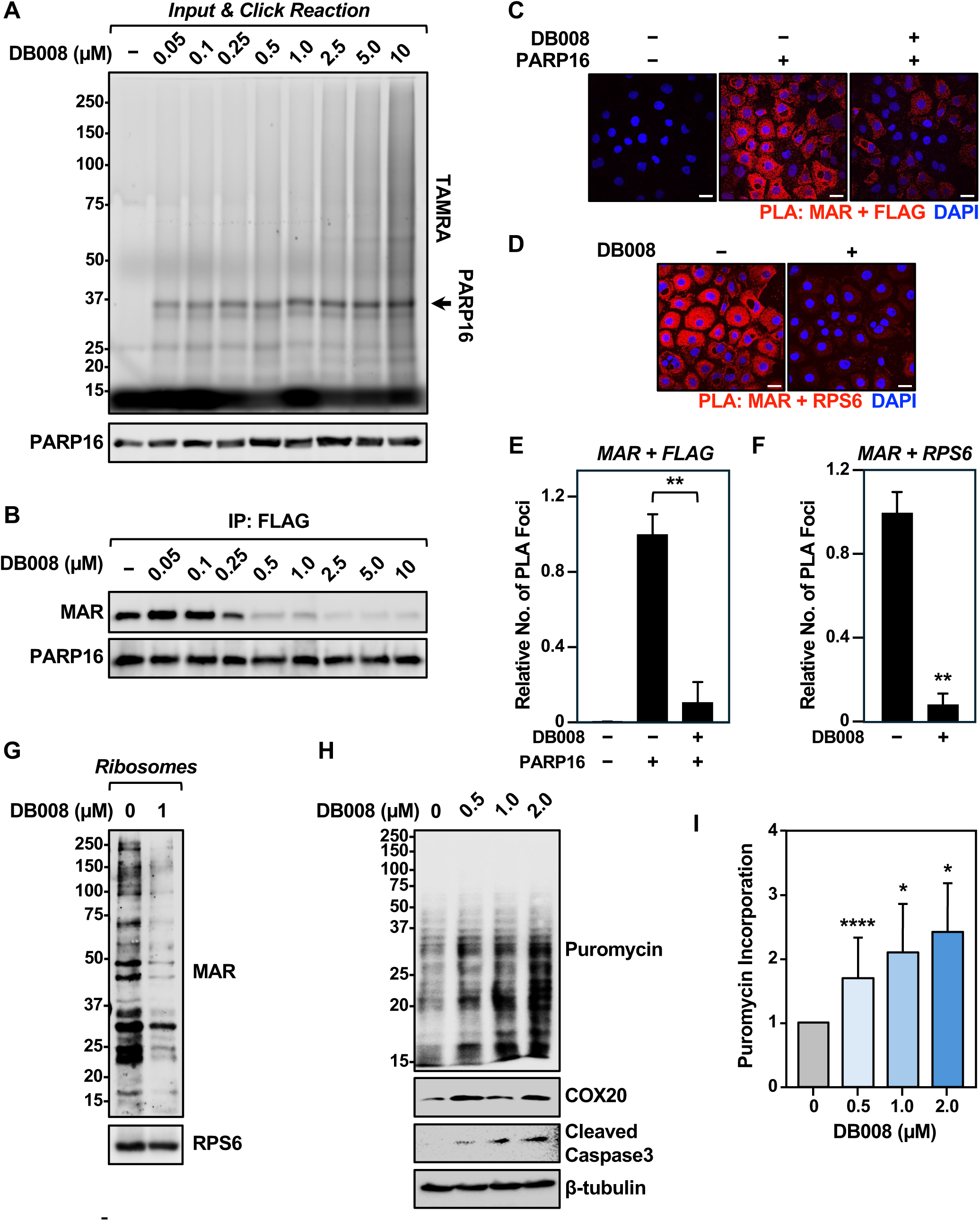
DB008 inhibits PARP16, decreases ribosome MARylation, and increases protein synthesis in ovarian cancer cells. **(A and B)** DB008 engages PARP16 and reduces its MARylation in ovarian cancer cells. OVCAR3 cells ectopically expressing FLAG-PARP16 were treated with 0 – 10 µM DB008 for 4 hours. (A) 20 µg of lysate were subjected to click chemistry to visualize DB008 target engagement with in-gel fluorescence (arrow indicates expected band size for FLAG-PARP16). **(B)** FLAG-PARP16 was immunoprecipitated from the remaining lysate and its MARylation was assessed with Western blotting. **(C and D)** In situ detection of (C) PARP16 autoMARylation and (D) RPS6 transMARylation. PLA of FLAG and MAR in OVCAR3 cells subjected to Dox-induced expression of FLAG-PARP16 and treated with 0 or 1 µM DB008 for 16 hours (C), or RPS6 and MAR in OVCAR3 cells treated with 0 or 1 µM DB008 for 16 hours (D). DNA was stained with DAPI. Scale bar is 25 µm. **(E and F)** Quantification of PLA foci in panel C (E) and panel D (F). Each bar represents the mean + SEM; n = 3. Welch’s t-test; ** = p < 0.01. **(G)** DB008 reduces ribosome MARylation. OVCAR3 cells were treated with 0 or 1 µM DB008 for 16 hours. Ribosomes were fractionated by ultracentrifugation and ribosome MARylation was assessed by Western blotting. RPS6 was used as a loading control. **(H and I)** DB008 enhances protein synthesis. OVCAR3 cells were treated with 0 – 2 µM DB008 for 16 hours. The rate of protein synthesis was then assessed with a puromycin incorporation assay. (D) Western blot showing puromycin incorporation, COX20, and cleaved caspase-3. β-tubulin was used as a loading control. (E) Quantification of puromycin signal in D. Each bar represents the mean + SEM; n = 3. Bars marked with asterisks are significantly different from the control; ANOVA; * = p < 0.05, **** = p < 0.0001.

### Cellular effects of DB008 depend on PARP16

To validate PARP16 as the target of DB008 in cells, we generated PARP16-knockout (KO) OVCAR3 cells (Fig. 3A). Immunofluorescent staining using a MAR-specific detection reagent shows perinuclear staining for MAR in control KO cells, which is abrogated upon treatment with DB008 (Fig. 3B). KO of PARP16 also reduced the MAR signal, which was not further affected by treatment with DB008 (Fig. 3B). We then measured protein synthesis using puromycin incorporation. As shown previously, we observed an increase in protein translation with DB008 in control cells (Fig. 3C). This increase was not observed in PARP16 KO cells. Similarly, protein aggregation increased upon treatment with DB008 or PARP16 KO, with no further aggregation observed in PARP16 KO cells with DB008 treatment (Fig. 3D). Lastly, we observed a significant effect on cell proliferation, where DB008 significantly reduced the growth of control, but not PARP16 KO cells (Fig. 3E), and DB008 also reduced colony formation in soft agar (Fig. 3F). These results confirm that the effects on MARylation, translation, protein aggregation, and cell growth observed by DB008 treatment are mediated by PARP16 activity.

**Figure 3.**
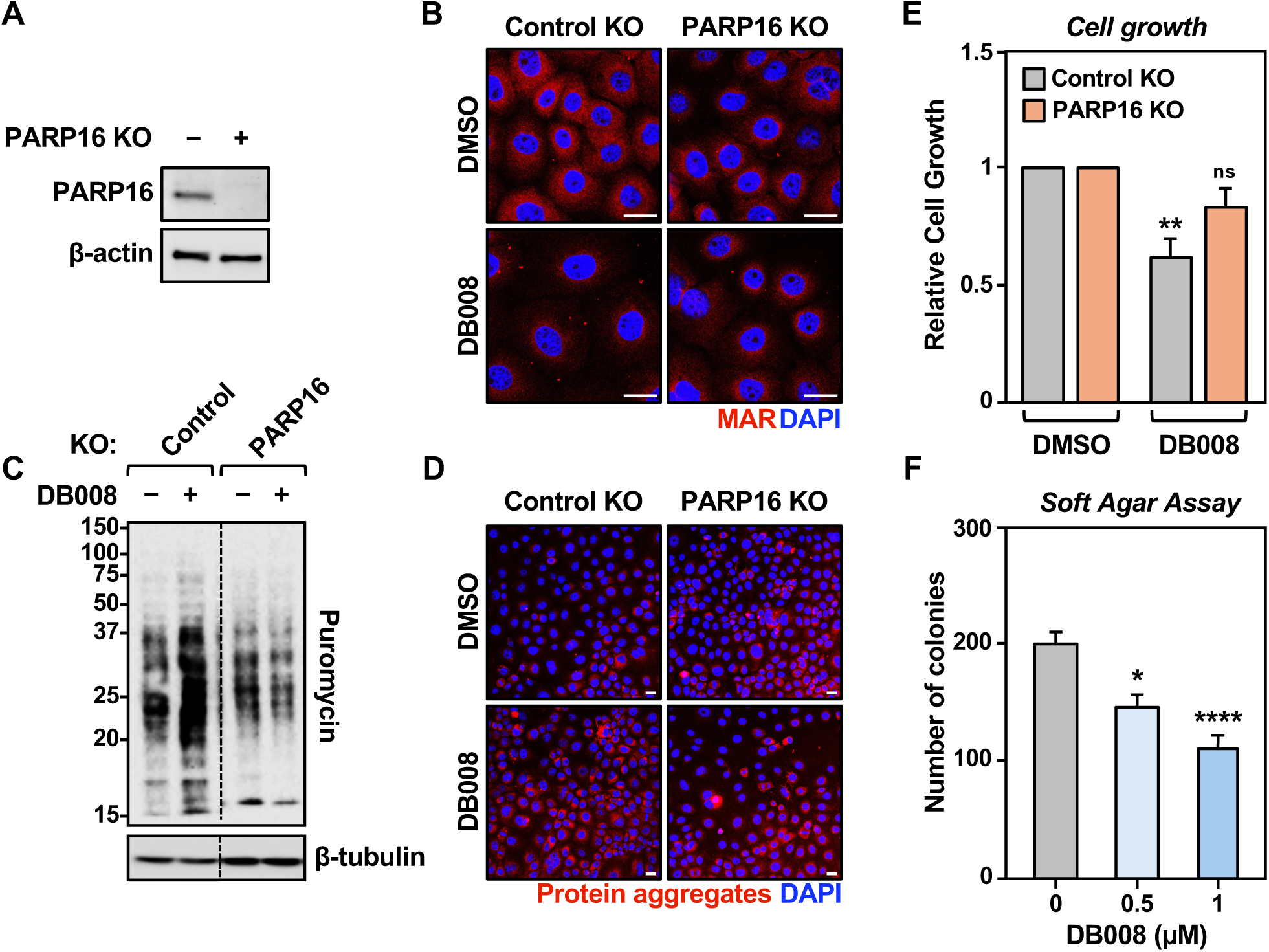
Cellular effects of DB008 depend on PARP16. **(A)** Western blot showing PARP16 protein expression is abolished in OVCAR3 cells with PARP16 KO. β-actin was used as a loading control. **(B)** Immunofluorescence with the MAR detection reagent in OVCAR3 Control or PARP16 KO cells treated with 0 or 1 µM DB008 for 16 hours. The DNA was stained with DAPI. Scale bar = 25 µm. **(C)** The DB008-dependent increase in protein synthesis requires PARP16 expression. OVCAR3 Control or PARP16 KO cells were treated with 0 or 1 µM DB008 for 16 hours. Protein synthesis was then assessed with a puromycin incorporation assay. β-tubulin was used as a loading control. **(D)** The DB008-dependent increase in protein aggregation requires PARP16 expression. OVCAR3 Control or PARP16 KO cells were treated with 0 or 1 µM DB008 for 16 hours. Protein aggregation was then assessed with the proteostat aggresome detection reagent. The DNA was stained with DAPI. Scale bar = 25 µm. **(E)** DB008 reduces growth of control but not OVCAR3 PARP16 KO cells. Cells were treated with 0 or 1 µM DB008 for 1 week. Cell growth was then determined with crystal violet staining. Each bar represents the mean + SEM; n = 3. Bars marked with asterisks are significantly different from the control; ANOVA; ** = p < 0.005; ns indicates no significance. **(F)** DB008 inhibits the anchorage-independent growth of ovarian cancer cells. OVCAR3 cells were grown in soft agar with media containing 0 – 1 µM DB008 for 3 weeks, replenishing media once per week. Anchorage-independent growth was then measured with crystal violet staining. Each bar represents the mean + SEM; n = 3. Bars marked with asterisks are significantly different from the control; ANOVA; * = p < 0.05, **** = p < 0.0001.

### Cells expressing PARP16 C169S are resistant to the effects of DB008

DB008 forms a covalent bond with Cys169 of PARP16 (20). To further validate PARP16 as the target of DB008 in cells, we generated OVCAR3 PARP16 KO cells ectopically expressing a FLAG-PARP16 mutant, where Cys169 was mutated to Ser (C169S), thereby disrupting the interaction. Immunoprecipitation of FLAG-PARP16 WT after treatment with DB008 showed a significant decrease in MARylation compared to control (Figs. 4, A and B). However, this reduction was not observed with FLAG-PARP16 C169S. No change in PARylation levels were evident (Fig. 4A). We examined anchorage-independent growth and found that OVCAR3 PARP16 KO cells stably expressing FLAG-PARP16 WT had a reduced ability for colony formation when treated with DB008 compared to control, while FLAG-PARP16 C169S cells were not affected by DB008 treatment (Fig. 4C). Similarly, increased protein synthesis (Figs. 4, D and E) and protein aggregation (Fig. 4F) were observed in FLAG-PARP16 WT expressing cells when treated with DB008 compared to control, but not in FLAG-PARP16 C169S expressing cells. Together, these results confirm that DB008 interacts with PARP16 through covalent bonding at C169, and this site is necessary for the biological effects observed by DB008-mediated inhibition of PARP16.

**Figure 4.**
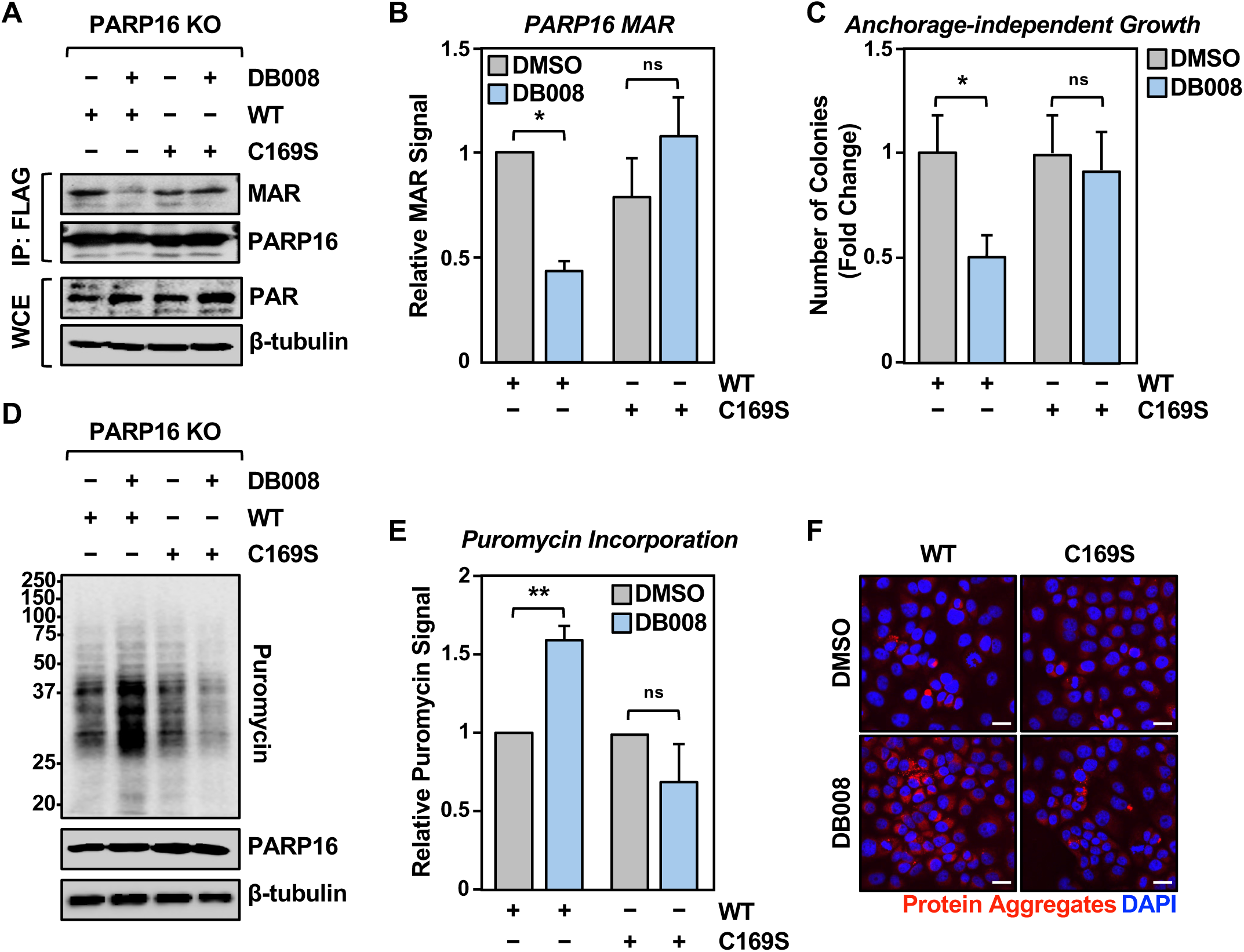
Cells expressing PARP16 C169S are resistant to the effects of DB008. **(A and B)** Inhibition by DB008 requires PARP16 C169. (A) IP-Western showing PARP16 MARylation (*top panel*) or PARylation levels in whole cell extracts (WCE; *bottom panel*) of OVCAR3 PARP16 KO cells stably expressing FLAG-PARP16 or FLAG-PARP16 C169S after treatment with 1 µM DB008 for 4 hours. (B) Quantification of MAR signal from A. Each bar represents the mean + SEM; n = 3. Bars marked with asterisks are significantly different from the control; ANOVA; * = p < 0.05; ns indicates no significance. **(C)** Anchorage-independent growth is unaffected in cells expressing PARP16 C169S. OVCAR3 PARP16 KO cells stably expressing PARP16 WT or PARP16 C169S were grown in soft agar with media containing 0 or 1 µM DB008 for 3 weeks, replenishing media once per week. Anchorage-independent growth was then measured with crystal violet staining. Quantification of crystal violet absorbance in B. Each bar represents the mean + SEM; n = 3. Bars marked with asterisks are significantly different from the control; ANOVA; * = p < 0.05; ns indicates no significance. **(D and E)** The DB008-dependent increase in protein synthesis requires PARP16 C169. (D) OVCAR3 PARP16 KO cells stably expressing PARP16 WT or PARP16 C169S were treated with 0 or 1 µM DB008 for 16 hours. Protein synthesis was then assessed with a puromycin incorporation assay. β-tubulin was used as a loading control. (E) Quantification of puromycin signal in D. Each bar represents the mean + SEM; n = 3. Bars marked with asterisks are significantly different from the control; ANOVA; ** = p < 0.01; ns indicates no significance. **(F)** The DB008-dependent increase in protein aggregation requires PARP16 C169. OVCAR3 PARP16 KO cells stably expressing PARP16 WT or PARP16 C169S were treated with 0 or 1 µM DB008 for 16 hours. Protein aggregation was then assessed with the proteostat aggresome detection reagent. The DNA was stained with DAPI. Scale bar = 25 µm.

### DB008 reduced ovarian cancer growth in vivo

We previously showed that genetic depletion of PARP16 inhibits the in vivo growth of ovarian cancer xenograft tumors (10). To test whether DB008 could recapitulate these results, we generated OVCAR3 xenograft tumors. Mice were randomized into vehicle or DB008 (50mg/kg) treatment groups (Fig. 5A). We observed significantly reduced tumor growth with DB008 treatment compared to vehicle (Figs. 5, B-D). Importantly we were able to detect targeted inhibition of PARP16 by fluorescent labeling of the DB008-PARP16 adduct with click chemistry directly from xenograft tumor lysates. These results indicate that pharmacologic inhibition of PARP16 with DB008 suppresses ovarian tumor growth in vivo through direct target engagement, providing proof-of-concept that PARP16 is a therapeutically actionable vulnerability in ovarian cancer.

**Figure 5.**
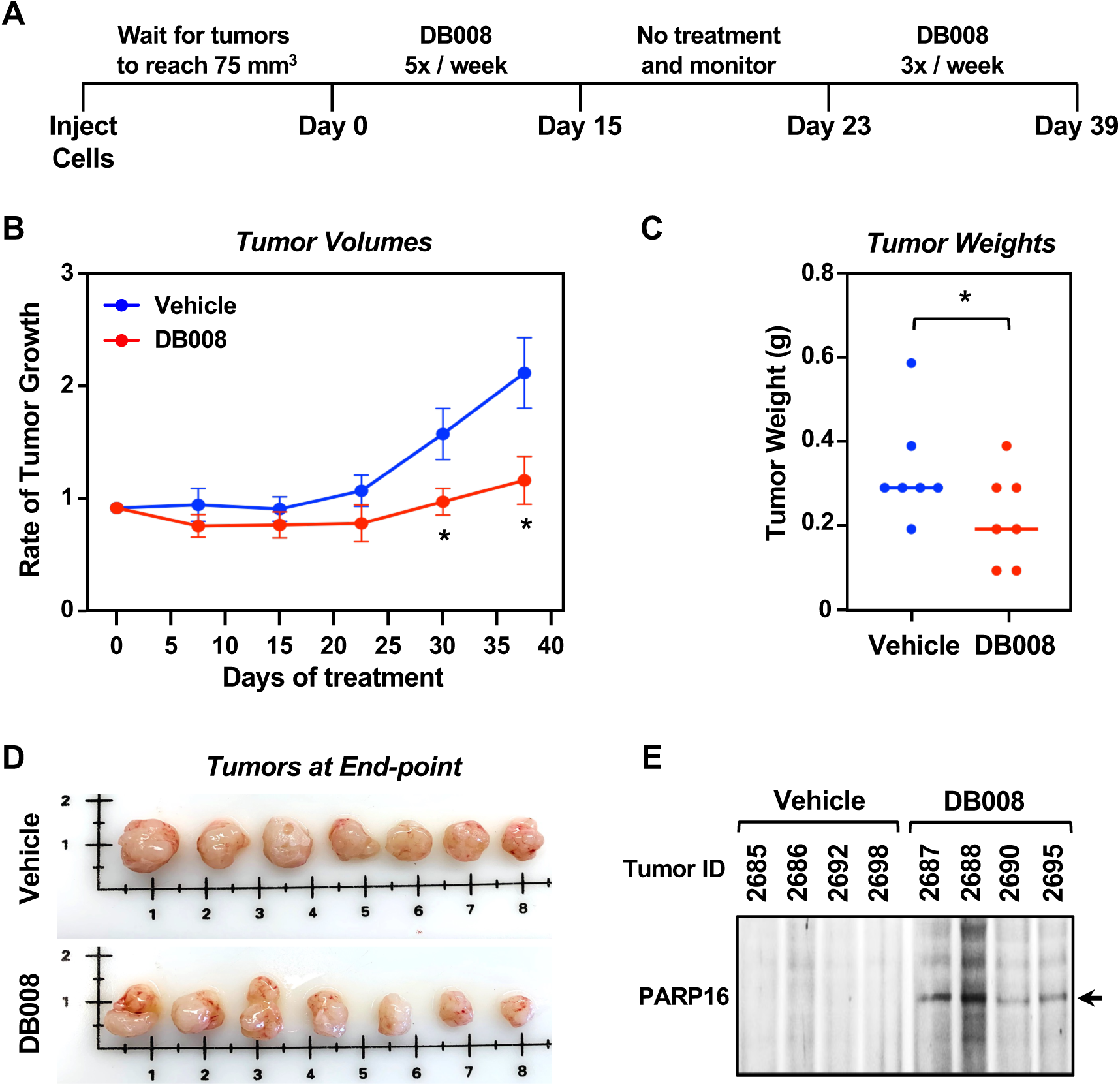
DB008 reduces ovarian cancer growth *in vivo*. **(A)** Schematic of the experimental timeline for the xenograft study. DB008 (50 mg/kg) or vehicle (saline) was administered by intraperitoneal injection 5 times per week for two weeks, monitored for one week, then treated 3 times per week for two weeks. **(B)** DB008 inhibits the growth of xenograft tumors formed from OVCAR3 cells. Graph showing the rate of tumor growth. Each point represents the mean ± SEM; n = 7. Points marked with asterisks are significantly different from the control; ANOVA; * = p < 0.05. **(C and D)** Tumor weights and images. (C) Weights of OVCAR3 xenograft tumors treated with Vehicle or DB008 at the end of the experiment (D) and images of OVCAR3 xenograft tumors treated with Vehicle or DB008 at the end of the experiment. The horizontal bar is the mean, and each point is an individual tumor; n = 7; ANOVA; * = p < 0.05. **(E)** DB008 inhibits PARP16 *in vivo.* In-gel fluorescence showing covalent inhibition of PARP16 by DB008 following fluorescent labeling of the DB008-PARP16 adduct with click chemistry in xenograft tumor lysates.

## Discussion

In this study, we established PARP16 as a druggable regulator of protein homeostasis in ovarian cancer and characterized DB008 as a potent, selective, and covalent inhibitor of its catalytic activity in this pathway. Our results demonstrate that pharmacologic inhibition of PARP16 phenocopies genetic depletion, leading to a disruption of ribosome MARylation, unregulated increases in protein synthesis, and the accumulation of toxic protein aggregates. This mechanism effectively suppresses ovarian cancer cell growth both in vitro and in vivo, providing a pharmacologic proof-of-concept for targeting this metabolic vulnerability.

### The role of PARP16 in protein homeostasis

Rapidly proliferating cancer cells require a precise balance between high-rate protein synthesis and stringent quality control. Our findings suggest that PARP16-mediated transMARylation of ribosomal proteins, such as RPS6, serves as a critical "brake" that fine-tunes translational output. By attenuating translation, PARP16 ensures that the cellular proteostasis and protein folding machinery are not overwhelmed (10). When this regulation is disrupted by DB008, the resulting surge in protein synthesis likely exceeds the capacity of cytosolic chaperones, leading to the proteotoxic stress and increased cleaved caspase-3 levels we observed. While some ribosomal proteins (e.g., RPL24) have been characterized as site-specific substrates for PARP16-mediated MARylation (10), this is an ongoing area of investigation in its infancy. Tools such as DB008 and NAD^+^ analogs will be useful for exploring specific sites of PARP16-mediated MARylation in future studies.

### Target engagement and specificity of DB008

A key challenge in the development of PARP inhibitors is achieving selectivity among the 17 family members. Previous work developed DB008, the first cysteine-targeted PARP inhibitor, which achieves selectivity through an ethynyl group at the C-6 position and covalent modification of a non-conserved cysteine in the PARP16 active site (20). We confirmed that DB008 has selectivity for PARP16 over PARP1 in vitro (IC_50_ 650 nM vs. 2.7 µM), and good proteome-wide selectivity in cells (Fig. 2A). We extend this work by showing for the first time that DB008 inhibits PARP16 MARylation in cells. Multiple assays demonstrated a reduction in PARP16 autoMARylation and transMARylation of ribosomal proteins upon DB008 treatment. Reduced ribosome MARylation led to anti-cancer effects, consistent with our previous results using PARP16 knockdown(10). Furthermore, our use of a C169S catalytic mutant provided definitive evidence that DB008 exerts its effects through direct covalent modification of PARP16. Cells expressing the C169S mutant were resistant to DB008-induced protein aggregation and growth inhibition, confirming that the compound’s anti-tumor activity is on-target.

### In vivo efficacy and clinical implications

Traditional chemotherapy remains the standard of care for ovarian cancer, but personalized treatments based on subtype and genetic risk factors are under investigation. The clearest examples are PARP1 inhibitors, which have considerably altered outcomes of some women with homologous-recombination deficient tumors (23). Other genetic alterations, such as *TP53* mutations and *CCNE1* amplification, are known to contribute to tumorigenesis and chemoresistance, but is unclear whether this knowledge can be translated into targeted therapies (24). Here, we investigated another signature alteration, *NMNAT2* amplification, which is characterized by high levels of cytosolic MARylation and is targetable through inhibition of PARP16. This approach is supported by a mechanistic understanding of the pathway that links cytosolic NAD^+^ synthesis by NMNAT-2 to PARP16-mediated transMARylation of ribosomes and translational regulation (10). We anticipate that inhibition of NMNAT-2 will produce similar effects, providing another actionable target and an opportunity for combination treatments that circumvent drug resistance. Future work will determine whether biomarkers such as *NMNAT2* expression and cytosolic MARylation predict responses to PARP16 inhibition in patient-derived models.

The ability of DB008 to significantly reduce tumor growth in OVCAR3 xenografts highlights the therapeutic potential of this strategy. Importantly, we detected direct target engagement within the xenograft tissues using click chemistry, linking the observed anti-tumor effects to the inhibition of PARP16. Given that many ovarian cancers rely on cytosolic NAD□ synthesis to support growth, PARP16 inhibitors like DB008 could represent a new class of targeted therapies that exploit the unique metabolic and translational requirements of these tumors. We note that the NMNAT-2/PARP16 pathway mediates similar effects in neuroblastoma cells and that NMNAT-2 overexpression promotes glioma growth, suggesting this therapeutic strategy may be applicable to other cancer types that share alterations in *NMNAT2* (10, 25, 26).

Finally, new studies are beginning to reveal a role of PARP16 as a target for non-oncological diseases. Apart from its functions in ovarian cancer, PARP16 also regulates protein homeostasis in normal physiology (27, 28). Recent studies have demonstrated that loss of PARP16 expression protects neurons from cell death in models of stroke and Alzheimer’s (29, 30). It will be of interest to determine whether small molecule inhibition of PARP16 is also protective in models of neurological disease.

## Experimental Procedures

### Antibodies, chemicals, and specialized reagents

The custom rabbit polyclonal antiserum against PARP16 was generated by Pocono Rabbit Farm and Laboratory by using a purified recombinant antigen comprising residues 1–279 with chromatography. The custom rabbit polyclonal antiserum against PARP1 was generated in-house by using purified recombinant amino-terminal half of PARP1 as an antigen (available from Active Motif, 39559; RRID:AB_2793257). The anti-mono(ADP-ribose) binding reagent (anti-MAR) and anti-poly(ADP-ribose) binding reagent (anti-PAR) were generated and purified in-house (available from Millipore Sigma, MABE1076; RRID: AB_2665469 and MABE1031; RRID:AB_2665467) (Gibson et al., 2017).

The other antibodies used were as follows: PARP16 (Abcam, ab84641; RRID:AB_1925296), β-tubulin (Abcam, ab6046; RRID:AB_2210370), β-Actin (Cell Signaling, 3700; RRID:AB-2442334), Puromycin (Millipore Sigma, MABE343; RRID:AB_2566826), FLAG (Sigma-Aldrich, F3165; RRID:AB_259529), COX20 (Proteintech, 25752-1-AP; RRID:AB_2880224), cleaved caspase-3 (Cell Signaling, 9661S; RRID:AB_2341188), rabbit IgG (Invitrogen, 10500C; RRID:AB_2532981), RPS6 (Cell Signaling, 2317S; RRIB:AB_2238583), Mono-ADP-ribose (Bio-Rad, TZA020), goat anti-rabbit HRP-conjugated IgG (ThermoFisher, 31460; RRID:AB_228341), goat anti-mouse HRP-conjugated IgG (ThermoFisher, 31430; RRID:AB_228307), BiSpyCatcher2-HRP (Bio-Rad, TZC002P), Alexa Fluor 594 donkey anti-rabbit IgG (ThermoFisher, A-21207; RRID:AB_141637), and Alexa Fluor 488 goat anti-mouse IgG (ThermoFisher, A-11001; RRID:AB_2534069).

For PARP16 inhibition, we used DB008 (MedChemExpress, HY-150221) (20). Other chemicals included: β-nicotinamide mononucleotide (NMN) (Sigma, N350), adenosine triphosphate (ThermoFisher, R0441), β-nicotinamide adenine dinucleotide (NAD^+^) (Sigma, N7381), Olaparib (MedChemExpress, HY-10162), PJ-34 (Abcam, ab120981), ADP-HPD (Sigma, 118415), and PDD 00017273 (MedChemExpress, HY-108360).

### Cell culture and treatments

#### Cell culture

Ovarian cancer OVCAR3 cells were purchased from the American Type Cell Culture (ATCC, HTB-161). The ovarian cancer cells were maintained in RPMI (Sigma, R8758) supplemented with 10% fetal bovine serum, 1% GlutaMax (ThermoFisher, 35050061) and 1% penicillin/streptomycin. OVCAR3 cells were grown for up to three months before being replaced with a low-passage cryogenic cell stock. Cell line was authenticated for cell type identity using the GenePrint 24 system (Promega, B1870), and confirmed as *Mycoplasma*-free every 6 months using the Universal Mycoplasma Detection Kit (ATCC, 30-1012K).

#### Cell treatments

For inhibition of PARP16, OVCAR3 cells were treated with the indicated concentrations of DB008 (0.5–2 µM) for 4 or 16 hours.

### Cloning and plasmid construction for PARP16 expression

#### Plasmid vectors for expression in bacteria

For GST-PARP16 and 10xHis-NMNAT-2, cDNA encoding human *Parp16* and *Nmnat2* were amplified by PCR, digested with restriction enzymes and ligated into pGEX-4T-1 (Sigma, GE28-9545-49) and pET-19b (Sigma, 69677-M) vectors, respectively. To make the antigen for the PARP16 antibody, cDNA encoding *Parp16* (1–279) was amplified by PCR, then inserted into pET-19b with Gibson assembly.

#### Lentiviral vectors for expression in mammalian cells

A plasmid expressing N-terminally tagged FLAG-PARP16 was generated using the cDNA for *Parp16* that was amplified from the pGEM-T cDNA clone obtained from Sino Biological (HG14445-G) and subcloned into the pCDH vector (System Biosciences, CD500B-1) as described previously (Challa et al., 2021). To generate a DB008-insensitive mutant, the C169S mutation was introduced into the pCDH-PARP16 plasmid using the QuikChange site-directed mutagenesis kit (Agilent, 200519).

#### Primers used for cloning

The following cloning primer sequences were used:

Cloning full length *Parp16* into pGEX-4T-1:

- *Parp16* Forward:

5’- GCGCGATCGCCATGCAGCCCTCAGGC -3’
- *Parp16* Reverse:

5’- GCGTTTAAACTTATCTTTTCGCACGATTCC -3’

Cloning *NMNAT2* into pET-19b:

- *NMNAT2* Forward:

5’- CTTTAAGAAGGAGATATACATGCATCATCATCATCATCATAC -3’
- *NMNAT2* Reverse:

5’- TTAGCAGCCGGATCCCTAGCCGGAGGCATTGATG -3’

Cloning *Parp16* (1–279) into pET-19b

- *Parp16* Forward:

5’-ACGACGACGACAAGCATATGCAGCCCTCAGGCTGGGCGGCCGCCAGGGAGGCGGCGGGCCGCGACATGCTGGCCGCCGACCTCCGGTGCAGCCTCTTCGCCTCGGCCCTGCAGAGCTACA -3’
- *Parp16* Reverse:

5’- TCGGGCTTTGTTAGCAGCCGGATCCTTAAGCCCTCTTGGGTGGCTTC -3’

Cloning full length *Parp16* into pCDH

- *Parp16* Forward:

5’- CTACTCTAGAGCTAGCGATGGACTACAAAGACGATGACGACAAGATGCAG CCCTCAGGCTGGGCGGCCGCCAGGGAGGCGGCGGGCCGCGACATGCTGGCC GCCGACCTCCGGTGCAGCC -3’
- *Parp16* Reverse:

5’- GGGAGAGGGGCGCGGCCGCGTTATCTTTTCGCACGATTCCAAAAGTGTTG -3’

Site-directed mutagenesis primers for *Parp16* in pCDH:

- *Parp16* C169S Forward:

5’- ACAATGGCCTGCACAGCCATCTGAACAAG -3’
- *Parp16* C169S Reverse:

5’- CTTGTTCAGATGGCTGTGCAGGCCATTGT -3’

### CRISPR/Cas9 genome editing

We used the Dharmacon CRISPR Design Tool to design two optimized single guide RNA (sgRNA) sequences for *Parp16*. Synthetic guide RNA was combined with TrueCut Cas9 Protein (ThermoFisher, A36498) and delivered into OVCAR3 cells with the Lipofectamine CRISPRMAX Transfection Reagent (ThermoFisher, CMAX00003).

#### sgRNA sequences

The following sgRNA target sequences were used:

- Parp16-1 (target in exon 1): 5’- AGAGCTACAAGCGCGACTCG -3’
- Parp16-2 (target in exon 1): 5’- TGAACGCACGCTCGCCCTTG -3’

### Expression and purification of recombinant PARP16 antigen, and generation of a polyclonal antiserum

For purification of the 10xHis-PARP16 (1–279) antigen, transformed BL21(DE3) cells were grown in Luria-Bertani medium at 37°C to an OD600 of 0.2, induced with 0.2 mM isopropyl-d-thiogalactoside (IPTG), then grown for 16 hours at 16°C. Bacteria were pelleted and lysed with lysis buffer (50 mM Na_2_HPO_4_ pH 4.0, 0.3 M NaCl, 1 mM PMSF, 8 M Urea). Lysate was sonicated, centrifuged to clear debris, and supernatant was incubated with Ni-NTA agarose (Qiagen, 30210). After incubation, beads were washed 3 times with wash buffer (50 mM Na_2_HPO_4_ pH 8.0, 0.5 M NaCl, and 8 M Urea), then washed 2 times with wash buffer containing 10 mM imidazole. After washing, PARP16 was eluted by incubating for 3 minutes with elution buffer (20 mM Tris pH 7.5, 100 mM NaCl, 8 M Urea, and 250 mM imidazole). Purified PARP16 (1–279) was aliquoted and flash frozen in liquid nitrogen. The antigen was sent to Pocono Rabbit Farm and Laboratory to generate the custom rabbit polyclonal antiserum against PARP16 (1–279).

### Expression and purification of recombinant full-length PARP16 and NMNAT-2

For purifications of GST-PARP16 and 10xHis-NMNAT-2, transformed BL21(DE3) cells were grown in Luria-Bertani medium at 37°C to an OD600 of 0.8, induced with 0.5 mM isopropyl-d-thiogalactoside (IPTG), then grown overnight at 18°C. Cells were harvested, resuspended GST Purification Buffer (50 mM HEPES pH 7.5, 0.5 M NaCl, 5 mM DTT, 10% glycerol, 1x protease inhibitor cocktail) or Ni-NTA Purification Buffer (25 mM HEPES pH 7.5, 0.5 M NaCl, 20 mM imidazole, 0.5 mM DTT, 10% glycerol, 1% NP-40, 1x protease inhibitor cocktail) for GST-PARP16 and 10xHis-NMNAT-2, respectively. Resuspended cells were flash-frozen in liquid nitrogen and stored at −80°C until purification. Cells were sonicated, then lysates were clarified with 40,000 x *g* for 30 minutes at 4°C. Purification was accomplished with Glutathione Sepharose 4B resin (ThermoFisher, 16100) and Ni-NTA agarose (Qiagen, 30210) for GST-PARP16 and 10xHis-NMNAT-2, respectively. Resins were washed with lysis buffer, then bound proteins were eluted with lysis buffer containing 10 mM reduced glutathione or 250 mM imidazole for GST-PARP16 and 10xHis-NMNAT-2, respectively. Eluted proteins were dialyzed overnight in Dialysis Buffer (10 mM HEPES pH 7.5, 200 mM NaCl, 0.5 mM DTT, 5% glycerol, 0.2% NP-40). Protein concentrations were estimated with the Bio-Rad Protein Assay Dye Reagent (Bio-Rad, 5000006). Single-use aliquots were flash-frozen in liquid nitrogen and stored at −80°C.

### In vitro autoMARylation assay

For the in vitro autoMARylation assay, purified GST-PARP16 (1 µM) and/or 10xHis-NMNAT-2 (1 µM) were combined with NAD^+^ or NMN/ATP (0 to 2 mM) in PARP Reaction Buffer (10 mM HEPES pH 7.2, 20 mM KCl, 10 mM MgCl_2_, 1 mM EGTA, 0.02% Tween-20, 0.5 mM DTT) in a reaction volume totaling 20 µL. The reaction was incubated at 25°C, 750 rpm for 2 hours, then quenched with SDS-PAGE sample buffer. The reaction products were separated using a 4-8% SDS-PAGE gel, then transferred to a nitrocellulose membrane. The membrane was then blotted for MAR and PARP16 using the anti-mono-ADP-ribose SpyTag antibody conjugated to BiSpyCatcher-HRP and the PARP16 antibody as described below. The signals were quantified with densitometry and plotted in Prism 10. The Km value was calculated with a nonlinear regression (curve fit) using the Michaelis-Menten equation.

### Estimation of IC_50_ values

#### PARP16-DB008 IC_50_ value

For determination of the PARP16-DB008 IC_50_ value, FLAG-PARP16 was immunoprecipitated from OVCAR3 cells as described below. FLAG-PARP16 (200 nM) was pre-incubated on ice with DB008 (0–10 µM) in PARP Activity Buffer (50 mM HEPES pH 7.2, 150 mM NaCl, 5 mM MgCl_2_, 1 mM EGTA, 1 mM DTT, 0.1% NP-40) for 30 minutes. Reactions were initiated with NAD^+^ (500 µM) and incubated at 25°C for 1 hour, then quenched with SDS-PAGE sample buffer. The reaction products were separated using a 4-8% SDS-PAGE gel, then transferred to a nitrocellulose membrane. The membrane was then blotted for MAR and PARP16 using the anti-mono-ADP-ribose SpyTag antibody conjugated to BiSpyCatcher-HRP and the PARP16 antibody as described above. The signals were quantified with densitometry and plotted in Prism 10. The IC_50_ value was calculated with a nonlinear regression (curve fit) using the sigmoidal dose-response equation.

#### PARP1 IC_50_ values

For determination of PARP1 IC_50_ values with DB008 and Olaparib, FLAG-PARP1 was purified from Sf9 cells as described previously (Huang et al., 2020). FLAG-PARP1 (25 nM) was pre-incubated on ice with DB008 (0–10 µM) or Olaparib (0–2.5 µM) in PARP Activity Buffer (50 mM HEPES pH 7.2, 150 mM NaCl, 5 mM MgCl_2_, 1 mM EGTA, 1 mM DTT, 0.1% NP-40, 0.1 mg/mL sheared DNA) for 30 minutes. Reactions were initiated with NAD^+^ (3 µM) and incubated at 25°C for 30 minutes, then quenched with SDS-PAGE sample buffer. The reaction products were separated using a 4-8% SDS-PAGE gel, then transferred to a nitrocellulose membrane. The membrane was then blotted for PAR and PARP1 using the anti-PAR detection reagent and the PARP1 antibody as described above. The signals were quantified with densitometry and plotted in Prism 10. The IC_50_ values were calculated with a nonlinear regression (curve fit) using the sigmoidal dose-response equation.

### Detecting DB008 target engagement in lysates with click chemistry

OVCAR3 cells or xenograft tissues were pulverized in Lysis Buffer (50 mM HEPES pH 7.4, 150 mM NaCl, 1 mM MgCl_2_, 1% Triton X-100) containing 0.1 mM TCEP, 250 nM ADP-HPD, 10 μM PJ-34, 1x protease inhibitor cocktail (Roche, 11697498001) and phosphatase inhibitors (10 mM sodium fluoride, 2 mM sodium orthovanadate, and 10 mM β-glycerophosphate). For experiments in OVCAR3 cells, whole cell lysates were prepared as described above. Lysates were centrifuged at 20,000 x *g* for 10 minutes at 4°C in a microcentrifuge. Protein concentrations were measured using the Bio-Rad Protein Assay Dye Reagent (5000006). 50 µg of protein was mixed with click buffer [100 µM TBTA, 1 mM CuSO_4_, 100 µM TAMRA-azide (Click Chemistry Tools), 1 mM TCEP, 1% SDS] in 1x PBS. The click reaction was incubated in dark for 1 hour at room temperature. After the click reaction, SDS-loading dye was added and the samples were boiled at 95°C for 5 minutes. The samples were loaded on a 12% SDS-PAGE gel. After electrophoresis, the gel was rinsed with water and imaged using the Rhodamine filter in ChemiDoc Gel Imaging System.

### Lysate preparation and immunoblotting

Cells were cultured and treated as described above before the preparation of cell extracts.

#### Preparation of whole cell lysates

At the conclusion of the treatments, the cells were washed twice with ice-cold PBS and resuspended in Lysis Buffer (20 mM Tris-HCl pH 7.5, 150 mM NaCl, 1 mM EDTA, 1 mM EGTA, 0.1% NP-40, 1% sodium deoxycholate, 0.1% SDS) containing 1 mM DTT, 250 nM ADP-HPD (or 10 µM PDD), 10 mM PJ-34, 1x protease inhibitor cocktail (Sigma, 11697498001) and 1x phosphatase inhibitor cocktail (Sigma, 4906845001). The cells were resuspended in Lysis Buffer and incubated on ice for 15 minutes, vortexed for 30 seconds and then centrifuged at 20,000 x *g* for 15 minutes at 4°C in a microcentrifuge to remove the cell debris. Protein concentrations were measured using the Bio-Rad Protein Assay Dye Reagent (5000006) and volumes of lysates containing equal total amounts of protein were mixed with 1/4 volume of 4x SDS-PAGE Loading Solution (250 mM Tris pH 6.8, 40% glycerol, 0.04% Bromophenol Blue, 4% SDS) and boiled at 95°C for 10 minutes, except for lysates used for MAR and PAR detection, which were incubated at 25°C for 10 minutes immediately before PAGE.

#### Nuclear and cytoplasmic fractionation

For the nuclear and cytoplasmic fractions, cells were washed twice with ice-cold PBS and collected by scraping in Isotonic Buffer (10 mM Tris-HCl pH 7.5, 2 mM MgCl_2_, 3 mM CaCl_2_, 0.3 M sucrose, 1 mM DTT) containing 1x protease inhibitor cocktail, 1x phosphatase inhibitor cocktail, 250 nM ADP-HPD (or 10 µM PDD), and 10 μM PJ-34. To swell the cells, they were incubated in the Isotonic Buffer on ice or 15 minutes and then lysed by the addition of 0.6% NP-40 followed by vortexing for 10 seconds. The nuclei from the lysed cells were pelleted by centrifugation at 10,000 x *g* for 30 seconds. The supernatant was collected and used as the cytoplasmic fraction. Cytoplasmic fractions were input material for immunoprecipitation and immunoblotting where indicated.

#### Ribosome fractionation

Ribosomal fractions were isolated from the cells as described previously (Kim et al., 2019). Briefly, the cells were plated into 15 cm dishes at 60% confluence one day prior to the assay. The next day, cells were treated with 1 µM DB008 or DMSO for 16 hours. After 16 hours of treatment, the cells were washed three times with ice-cold PBS and scraped gently into 1.5 mL Buffer A (50 mM Tris-HCl pH 7.5, 250 mM sucrose, 250 mM KCl, 5 mM MgCl_2_) supplemented with protease inhibitors, phosphatase inhibitors, PARG inhibitor, and PJ-34 as described above. NP-40 was then added to a final concentration of 0.7% (v/v) and the cells were incubated on ice for 15 minutes with frequent mixing. Five percent of each lysate was removed for an input sample. The remaining portion of each lysate was centrifuged at 750 x *g* for 10 minutes at 4°C in a microcentrifuge, and the supernatants were centrifuged again at 12500 x *g* for 10 minutes at 4°C in a microcentrifuge to remove nuclear proteins and transferred to a new tube. The concentration of KCl in the lysates was adjusted to 500 mM using a 3 M KCl stock and the lysates were loaded onto a 2.5 mL sucrose cushion (50 mM Tris pH 7.5, 1 M sucrose, 0.5 M KCl, 5 mM MgCl_2_) in polypropylene tubes (Beckman Coulter, 328874). The samples were centrifuged for 4 hours at 210,000 x *g* in a Beckman coulter Optima L-80 XP ultracentrifuge using a SW60Ti rotor. After centrifugation, the supernatant and sucrose cushion in each tube were discarded, and the ribosomal pellet was resuspended in Buffer C (50 mM Tris pH 7.5, 25 mM KCl, 5 mM MgCl_2_) supplemented with protease, phosphatase, and PARG inhibitors. The protein concentration was estimated, and MAR visualized with immunoblotting using the anti-MAR detection reagent.

#### Immunoblotting

Protein lysates were run on SDS-PAGE gels and transferred to nitrocellulose membranes. After blocking with 5% nonfat milk or BSA in TBST, the membranes were incubated with the primary antibodies described above with 5% nonfat milk or BSA in TBST, followed by anti-rabbit HRP-conjugated IgG (1:2000) or anti-mouse HRP-conjugated IgG (1:2000). Immunoblot signals were captured using a luminol-based enhanced chemiluminescence (ECL) HRP substrate (SuperSignal™ West Pico; Thermo Scientific, 34580) or an ultra-sensitive enhanced chemiluminescence HRP substrate (SuperSignal™ West Femto; Thermo Scientific, 34094) and a ChemiDoc imaging system (Bio-Rad).

### Generation of cell lines for ectopic expression

Cells were transduced with lentivirus for ectopic expression. Lentiviruses were generated by transfection of the pCDH constructs described above, together with (i) an expression vector for the VSV-G envelope protein (pCMV-VSV-G, Addgene 8454; RRID: Addgene_8454), (ii) an expression vector for GAG-Pol-Rev (psPAX2, Addgene 12260; RRID: Addgene_12260), and (iii) a vector to aid with translation initiation (pAdVAntage, Promega E1711) into 293T cells with the GeneJuice transfection reagent (Novagen, 70967) according to the manufacturer’s protocol. The resulting viruses were used to infect OVCAR3 PARP16 KO cells in the presence of 7.5 µg/mL polybrene for 24 hours after initial 293T transfection. Stably transduced cells were selected with 1 µg/mL puromycin (Sigma, P9620).

### Immunoprecipitation of FLAG-PARP16

OVCAR3 cells were seeded at ∼2□×□10^6^ cells per 15□cm plate and transfected at ∼60% confluence with pCDH FLAG-PARP16 WT or C169S using Lipofectamine 3000 (Invitrogen, L3000015) for 16 hours according to the manufacturer’s protocol. Appropriate cell treatments were performed as described above. The cells were collected and whole cell extracts were prepared as described above. The resulting extracts were incubated with equilibrated anti-FLAG M2 beads (Sigma-Aldrich, A2220) for 1 or 16 hours at 4°C with gentle mixing. The beads were washed five times with gentle mixing for 10 minutes at 4°C with Immunoaffinity Purification Wash Buffer (25 mM Tris-HCl pH 7.5, 150 mM NaCl, 0.1% NP-40 and 1x complete protease inhibitor cocktail). FLAG-tagged proteins were then eluted by incubation with 0.1 mg/mL FLAG peptide in Immunoaffinity Purification Wash Buffer for 30 minutes at 4°C. The eluted immunoprecipitate was subjected to immunoblotting as described above.

### Proximity ligation assays (PLA)

PLAs were performed using a Duolink proximity ligation kit (Sigma, DUO92008) following the manufacturer’s instructions. Briefly, cells were plated on 8-well culture slides (Corning, 354108). At the end of treatments, the cells were washed once with PBS, fixed in 4% paraformaldehyde for 15 minutes at room temperature, and washed three times with PBS. The cells were permeabilized for 10 minutes at −20°C using ice-cold methanol, washed three times with PBS, and incubated in Duolink Blocking Solution for 1 hour at 37°C in a humidified chamber. Excess Blocking Solution was removed by tapping and the cells were incubated in the primary antibody pairs: mouse FLAG (1:500) or mouse RPS6 (1:100) and rabbit MAR detection reagent (20 µg/ml) antibodies in Duolink Antibody Diluent. The slides were incubated overnight at 4°C in a humidified chamber. Following the overnight incubation, the cells were washed twice for 5 minutes each with Wash Buffer A (10 mM Tris-HCl pH 7.4, 150 mM NaCl, and 0.05% Tween). The slides were incubated with PLUS and MINUS PLA probes 1:5 in the Duolink Antibody Diluent for 1h at 37°C in a humidified chamber, followed by two washes with Washing Buffer A for 5 minutes each. The slides were then incubated with Ligation Solution (1:40 dilution of the ligase in 1x Ligation Buffer) for 30 minutes at 37°C in a humidified chamber, followed by two washes with Wash Buffer A for 5 minutes each. The cells were then incubated in the Amplification Solution (1:80 dilution of the Polymerase in 1x Amplification Buffer) for 100 minutes at 37°C in a humidified chamber protected from light. The cells were washed twice with Wash Buffer B (200 mM Tris-HCl pH 7.5, 100 mM NaCl) for 10 minutes each followed by a wash with 0.01x Wash Buffer B for 2 minutes. The stained cells were mounted on microslides using VectaShield Antifade Mounting Medium (Vector laboratories, H-1200-10) with DAPI DNA stain. Images were acquired using an inverted Nikon Ti-2 confocal microscope using an immersion oil 60X objective at room temperature.

### Puromycin incorporation

Puromycin incorporation assays were performed as described previously (Schmidt et al., 2009). For DB008 treatment, cells were plated at 60% confluence in 6-well plates. The following day, cells were treated with 0.5–2 µM DB008 or DMSO for 16 hours. Cells were then treated with 10 μg/mL puromycin for 15 minutes at 37°C and whole cell extracts were prepared from these cells as described above. Puromycin incorporation was visualized by immunoblotting with an antibody against puromycin (Sigma, MABE343).

### Immunofluorescent staining of cultured cells

The following microscopy-based protocols for cultured cells were used to determine cellular MAR localization and protein aggregation levels in cells.

#### Immunostaining for MAR

OVCAR3 cells were seeded on 8-well chambered slides (ThermoFisher, 154534) one day prior to the experiment. The cells were washed twice with PBS, fixed in 4% paraformaldehyde for 15 minutes at room temperature, and washed three times with PBS. The cells were permeabilized for 5 minutes using Permeabilization Buffer (PBS containing 0.01% Triton X-100), washed three times with PBS, and incubated for 1 hour at room temperature in Blocking Solution (PBS containing 1% BSA, 10% FBS, 0.3 M Glycine and 0.1% Tween-20). The fixed cells were incubated with MAR detection reagents at a concentration of 20 μg/mL in PBS overnight at 4°C, followed by three washes with PBS. The cells were then incubated with Alexa Fluor 594 donkey anti-rabbit IgG (ThermoFisher, A-21207) and/or Alexa Fluor 488 goat anti-mouse IgG (ThermoFisher, A-11001) each at a 1:500 dilution in PBS for 1 hour at room temperature. After incubation, the cells were washed three times with PBS. Finally, coverslips were placed on cells coated with VectaShield Antifade Mounting Medium with DAPI (Vector Laboratories, H-1200) and images were acquired using an inverted Zeiss LSM 780 confocal microscope.

#### Protein aggregation assay

The levels of protein aggregation in cells were measured using the Proteostat Protein Aggregation Kit (Enzo, ENZ-51035) according to the manufacturer’s protocol. Briefly, the cells were plated in 8-well chambered slides and treated ± cycloheximide (10 μg/mL) in normal growth media for 16 hours. The cells were then fixed in 4% paraformaldehyde for 30 minutes, permeabilized in Permeabilization Buffer (1x assay buffer containing 0.5% Triton X-100, 3 mM EDTA pH 8.0) for 30 minutes at 4°C. The cells were then treated with the Proteostat aggresome detection reagent for 30 minutes at room temperature. After washing with PBS, the coverslips were fixed with VectaShield Antifade Mounting Medium with DAPI (Vector Laboratories, H-1200), and confocal imaging was performed using an inverted Zeiss LSM 780 confocal microscope.

#### Image analysis

The fluorescence intensities captured by the confocal imaging were analyzed with Fiji ImageJ (Schindelin et al., 2012). The intensity and contrast of the images were further adjusted in Microsoft PowerPoint. The same intensity and contrast were applied to all images in an experiment.

### Cell growth assays

OVCAR3 control and PARP16 knockout cells were plated at a density of 3,000 cells per well in a 96-well plate. The cells were then treated with either 1 µM DB008 or DMSO. The cells were fixed with 4% paraformaldehyde on 0 and 7 days after treatment and washed with water. The fixed cells were then stained with crystal violet solution (0.5% crystal violet in 20% methanol) for 30 minutes with gentle shaking at room temperature. The stained cells were washed with water and air dried. The crystal violet was then dissolved in 10% acetic acid and the absorbance at 570 nm was measured using a spectrophotometer. The absorbance of a blank well was subtracted from the samples and the values were normalized to the values at day 0. Three independent biological replicates were performed to ensure reproducibility. Statistical differences were determined using one-way ANOVA.

### Anchorage-independent growth assays

The bottom agar layer was made by adding 1 mL of 6% agar (Thermo Scientific, J10907-22) in complete growth medium in a 12-well plate. The agar layer was allowed to solidify at 37°C for 30 minutes. The top layer, 1 mL of 3% agar containing 10,000 cells, was added gently over the bottom layer and the cells were cultured for 3 weeks. The cells were replenished with 300 μL fresh medium once per week. After 3 weeks, the colonies were stained with crystal violet solution (0.5% crystal violet in 20% methanol). Excess crystal violet stain was washed with water and the plates were imaged. The number of colonies per well were manually counted. Three independent biological replicates were performed, and statistical differences were determined using a one-way ANOVA.

### Mouse xenograft studies

All animal experiments were performed in compliance with the Institutional Animal Care and Use Committee (IACUC) at the UT Southwestern Medical Center. Female NOD/SCID/gamma (NSG) mice at 6-8 weeks of age were used. To establish ovarian cancer xenografts, 2 × 10^6^ OVCAR3 cells were injected subcutaneously in 100 μL into the flank of the mice in a 1:1 ratio of PBS and Matrigel (Fisher, CB 40230). Mice were treated with DB008 (50 mg/kg) or vehicle (saline) by intraperitoneal injection 5 times per week for two weeks, monitored for one week, then treated 3 times per week for two weeks. Mouse weight was monitored once per week and tumor growth was measured using electronic calipers once per week. Tumor volumes were calculated using a modified ellipsoid formula: Tumor volume = ½ (length × width^2^). At the end of the experiment, mice were euthanized to collect the xenograft tissue. The tissue was cut into several small pieces, and separate portions were either snap-frozen in liquid nitrogen or fixed using 4% paraformaldehyde. The frozen tissues were pulverized using a tissue mill and lysed in Whole Cell Lysis Buffer (20 mM Tris-HCl pH 7.5, 150 mM NaCl, 1 mM EDTA, 1 mM EGTA, 1% NP-40, 1% sodium deoxycholate, 0.1% SDS, 1 mM DTT, 250 nM ADP-HPD, 10 μM PJ-34 supplemented with protease and phosphatase inhibitors) for protein extraction. The protein samples were analyzed by immunoblotting as described above.

### Quantification and statistical analyses

All biochemical and cellular assays were performed a minimum of two times with independent biological samples and analyzed by Image Lab 6.0. For the xenograft study, seven mice were in each treatment group. Statistical analyses were performed using GraphPad Prism 10. All tests and p values are provided in the corresponding figures or figure legends. In all figures, the p values are shown as: *, p < 0.05; **, p < 0.01; ***, p < 0.001, ****, p < 0.0001, or other values as specified.

## Data Availability

All data and reagents presented within this article are available upon request.

## Supporting Information

None.

## Acknowledgements and Funding

We thank Daniel S. Bejan and Michael S. Cohen for sharing the DB008 compound for our initial experiments. We also thank members of the Kraus lab for their helpful comments and support. This work was supported by grants from the NIH (R01 DK069710) and the U.S. Congressionally Directed Medical Research Program (CDMRP) Ovarian Cancer Research Program (OCRP) (OC200311, OC230196, OC250231) to W.L.K. and funds from the Cecil H. and Ida Green Center for Reproductive Biology Sciences Endowment to W.L.K.

## Author Contributions

S.C.- conceptualization, formal analysis, investigation, validation, visualization, data curation. M.D.- conceptualization, formal analysis, investigation, validation, visualization, writing- original draft preparation, writing- review and editing. J.C.A.- formal analysis, investigation. K.P.- formal analysis, investigation. L.Y.- formal analysis, investigation. C.V.C.-supervision, visualization, writing- original draft preparation, writing- review and editing. W.L.K.- conceptualization, funding acquisition, project administration, supervision, visualization, writing- review and editing.

